# Object Recognition in P14 mice

**DOI:** 10.1101/850933

**Authors:** Arvind Chandrakantan, Adam C. Adler, Fred A. Pereira

**Affiliations:** Texas Children’s Hospital; Baylor College of Medicine; Huffington Center of Aging

## Abstract

Object Recognition is a task which involves multiple brain areas for successful completion. This assay is non-invasive, is an enriched learning task, and relies upon on encoded memory for successful completion. In this study, we have demonstrated that neonatal mice can perform the task

## Introduction

There are many behavioral learning tests for adult mice, however, to date, there is a lack of methodological behavior learning testing for neonatal mice. Mice have unique visual development and generally do not open their eyes until their 12^th^ post gestational day (P12)[1]. Even after eyelid opening, and obtaining visual capability, there are several differences between mouse and human vision, namely: 1) sky versus ground optimization of vision is geared towards ground [2] [3, 4] [5]; and 2) predominance of rods over cones; and 3) a general visual system resembling primate/carnivore architecture[6]. In neonatal mice, the timing and sequence of visual inputs and cortical processing remain to be defined. What is known is that the rodent visual apparatus has high-level visual processing acumen with invariant object recognition [7, 8].

The object recognition test is typically performed after mice are weaned from their mother and in the stage of pre-adolescence with fully developed visual ability at P30 or later. This assay has several variations. The one used here was a one-trial learning task, wherein the animal displays a preference for a novel object over a familiar one [9, 10] after a single session of exposure to 2 similar objects. This activity relies on several areas of the brain, most notably: 1) the visual processing system, 2) the hippocampus, where new memories and experiences are encoded [11, 12] and 3) the medial temporal lobe [13, 14]. Sleep is important in consolidating memory related to this task[15] as sleep deprivation affects memory encoding[16].

It is unknown whether neonatal P14 mice have the capacity to perform a task which involve multisystem cuing, including visual recognition, of different laboratory objects very soon after eyelid opening (EO). The period around EO is characterized by a flood of synaptic activity involving numerous regions of the brain [17, 18] Therefore, neonatal mouse would require several faculties to perform the OR test successfully.

## Materials & Methods

The institutional animal care and use committee (IACUC) at Baylor College of Medicine has approved the use of animals in this study.

### Animals

All animals are from the C57BL6 wild type line (Jackson Laboratories). Animals were cared for in the NRI vivarium with access to food and water and checked daily. The day light cycle of the vivarium was 12h/12h with lights out at 8 PM and on at 8 AM. Animals were handled per protocol below. At the conclusion of the analysis, animals were euthanized utilizing CO_2_ per protocol. Animals were euthanized painlessly by approved IACUC methods and under the rules and regulations at Baylor College of Medicine. Specifically, we exposed the mice to compressed carbon dioxide (100%) obtained from cylinders. The chamber pre-filled to a 75% carbon dioxide concentration to induce unconsciousness and death more quickly. Our attending veterinarian and the IACUC have approved these procedures. All procedures are also consistent with the recommendation of the Panel on Euthanasia of the American Veterinary Medical Association. All investigators have completed all necessary pre-requisites and have been certified to work with rodents.

### Setup

The arena in which the test is performed is a small 22cm × 44cm plexiglass chamber, surrounded on three sides by a white screen to limit spatial information and prevent spatial biases that may influence object exploration. The total exploration time per session is 5 minutes with an intersession interval of 24 hours. An angled mirror is placed behind the cage to enable the observer to see the back side of the arena. Mice are placed into the arena and the time spent interacting with the objects is recorded (ANY-maze video tracking system (Stoelting Co., USA). On day # 2 one of the objects is changed with one of 3 validated objects (all internally validated as novel within our core facility) and the time spent interacting with each object is recorded.

### Protocol

The Object Recognition(OR) protocol utilized occurred over 2 contiguous days with an intersession interval (ISI) of 24 hours between testing sessions. Familiarization of the arena and object occurred on day 1 and testing on day 2 when the novel object is introduced. The first day of testing occurred on day P13 (both sides with the same object), and the second day with the novel object on P14, as calculated from P0, the day of birth. The time of testing was identical on both days, and the mice were all transferred in the same order as the prior day. The mice were habituated with the handler to avoid stress response. Specifically, a sleeved and gloved hand was placed in their cages for 1 minute each day for one week prior to the test being performed. NOR testing was done during the day. All of the pups were transferred from their home cage with their parents into a holding cage to habituate for 5 minutes. After the test, they were returned to their home cage with their parents. The objects were cleaned with alcohol and dried between animals to prevent cuing.The total test time was 5 minutes for each day and for each mouse, and the metrics were recorded as the number of interactions with each object, mean time spent interacting with each object, and the latency to the first-time object exploration. The novel object was rotated randomly on day 2 and both the object as well as laterality.

**Figure 1:**
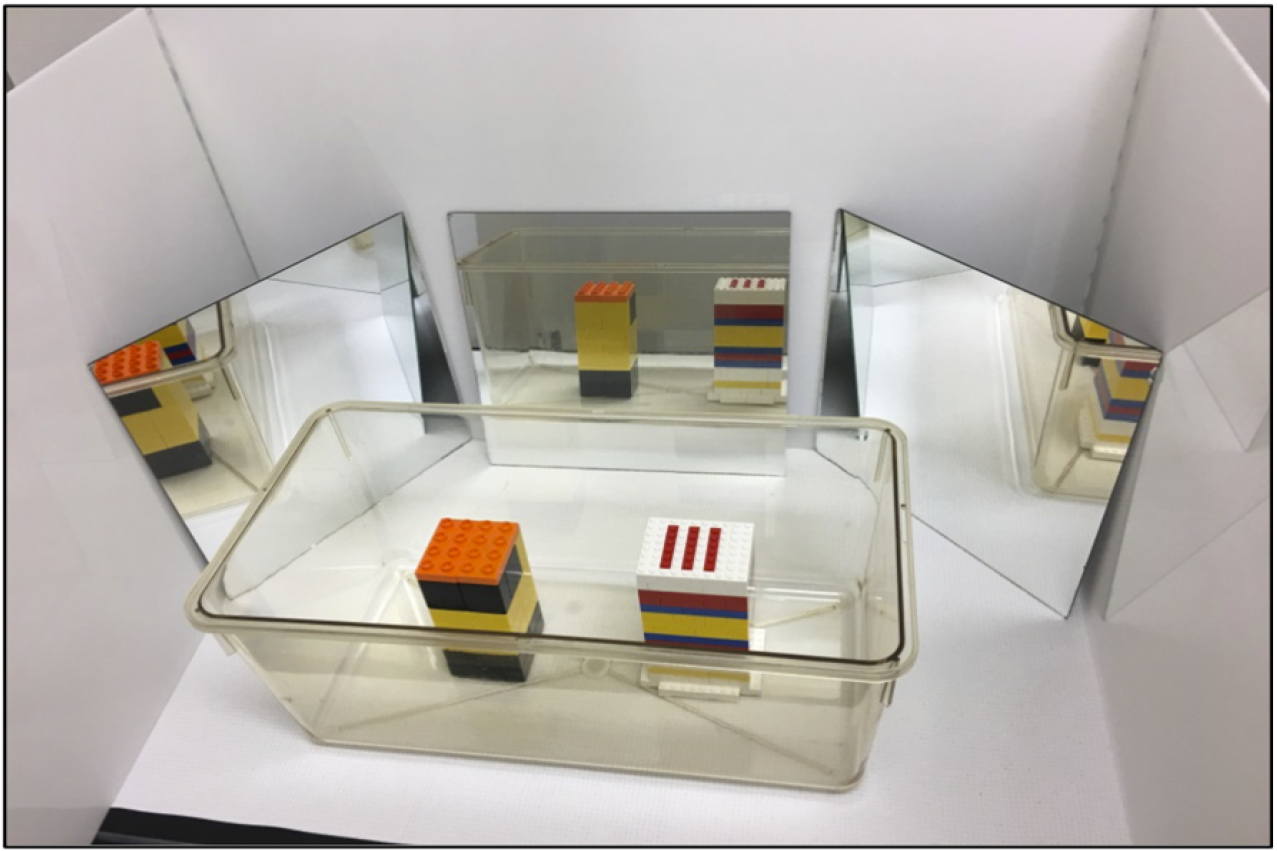
The NOR setup includes 3 offset mirrors to document activity behind the objects, as well as measured placement of the objects from the center of the container.

Exploratory behavior was defined as nose-directory behavior towards the object [19]. Non-nose directed behaviors including climbing and pawing of the objects without nasal cuing were excluded from the analysis. All analysis was done post-hoc in a blinded fashion. The scoring algorithm followed the protocol from Leger et al[20].

## Results

There were a total of 16 males and 15 females over the entire group. The number of interactions and time spent with each object within each batch of mice were compared using paired T-tests. The time spent and number of interactions between objects were not statistically significant on Day 1, where the same object was presented on both sides of the enclosure (see Supplemental data). On Day 2, when the novel object was placed, each group demonstrated statistically significant increases in both number of times interacted with, p-value (95% CI): <0.001(−15.1; −8.0), and time spent with the novel object, p-value (95% CI): <0.001(−9.6; −5.7). Of note, the number of interactions increased for both objects on Day 2 compared to Day 1, p-value (95% CI):0.003(−17.5; −5.9). This may have occurred because the mice had become familiar with the enclosure and had less anxiety about being closer to the center of the enclosure where the novel objects were placed. The increase in interactions with the novel object were statistically significant over the known object demonstrating the mice had the capacity to recognize object novelty. Furthermore, the discrimination ratio, which measures recognition memory sensitivity[21], was found to be statistically significant with regards to the novel object (Figure 2) p-value (95% CI) <.00001, (0.457, 0.73). The level of exploration is available in the Supplemental data.

**Figure 2.**
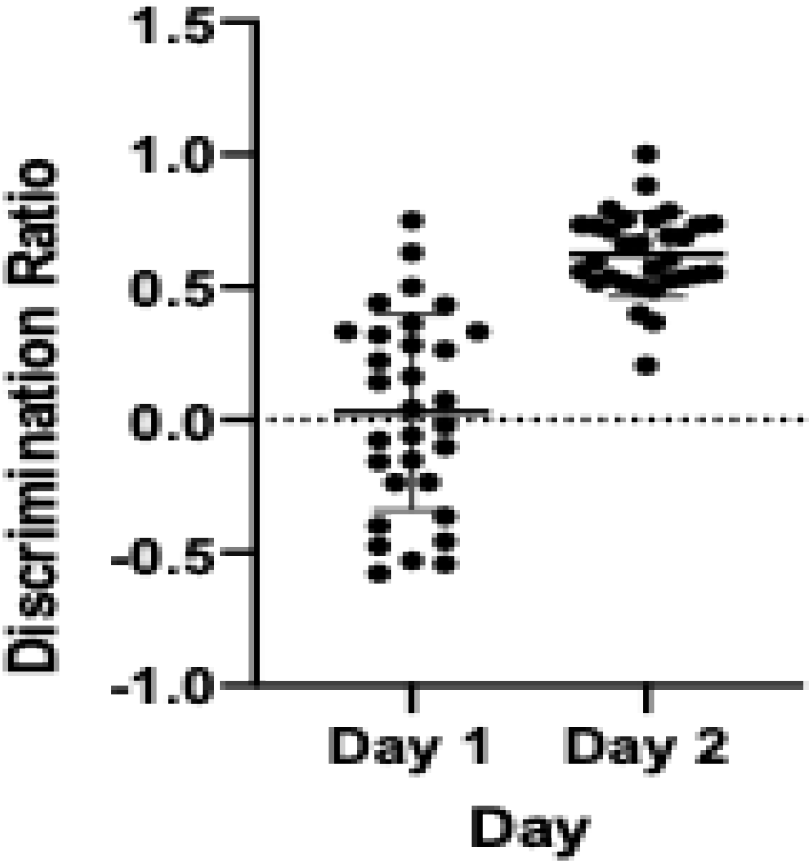
demonstrates the discrimination ratio between Day 1 and Day 2 adjusted for novel object placement on day 2, demonstrating recognition memory sensitivity without bias.

There was no statistically significance difference in weight between batches or between male and female mice, *p*=0.54 (0.59, 0.32) or with regards to exploration characteristics (number of interactions or time spent). The supplemental video file shows mouse exploration.

## Discussion

Mice have poor vision at young ages where there is a high incidence of myopia [22]. The majority of exploration occurs through direct contact, including climbing, pawing, sniffing, and other behaviors[10], which involve direct object interaction utilizing nasal direction for the NOR test. Importantly, if the animals were unable to recognize the novelty, then the interaction would be random between objects, and the discrimination ratio would be 0. All the mice within and across the different batches displayed an overt preference for the novel object, both in time spent and number of interactions. the OR assay represents a non-stressful, non-aversive means of interrogating encoded memories in young mice. This study demonstrates that the OR test is reproducible when used with neonatal mice as early as P14. when used with neonatal mice as early as P14.

Object recognition has been used as one of the primary learning and memory tasks to assess the effect of environmental stimuli on hippocampal function [23]. Object recognition interrogates several brain areas including the CA-1 region of the hippocampus [24]. This is evidenced by delayed and disrupted object memory especially with a delayed testing interval in mice with induced hippocampal deficits[25]. To ensure consistent interpretation of the OR testand to decrease variability in the task, the object type must be consistent for a robust interaction between the mice and the objects[26, 27]. Therefore, context, placing, and accurate postnatal timing are all important in executing the OR test correctly for neonatal mice. It is easily administered, quickly tests function, and can be manipulated based on experimental goals, as long as basic tenets of the test are adhered to[20]

## Conclusion

The OR test can be successfully performed in P14 neonatal mice, demonstrating that this is non-stressful, non-aversive examination can be used soon after eyelid opening (EO). Use of the OR test will inform future studies utilizing neonatal mice in interrogating early learning and memory.

## Supplemental Files

P14 Excel spreadsheet with gender, weight, and raw data of mice Videos of mouse motion indicating the mice are exploring the objects

